# *In situ* structures of secretins from bacterial type II secretion system reveal their membrane interactions and translocation process

**DOI:** 10.1101/2023.01.10.523476

**Authors:** Zhili Yu, Yaoming Wu, Muyuan Chen, Tong Huo, Wei Zheng, Steven J. Ludtke, Xiaodong Shi, Zhao Wang

## Abstract

The GspD secretin is the outer membrane channel of the bacterial type II secretion system (T2SS) which secrets diverse effector proteins or toxins that cause severe diseases such as diarrhea and cholera. GspD needs to translocate from the inner to the outer membrane to exert its function, and this process is an essential step for T2SS to assemble. Here, we investigate two types of secretins discovered so far in *Escherichia coli*, GspD_α_ and GspD_β_, respectively. By electron cryotomography subtomogram averaging, we determine *in situ* structures of all the key intermediate states of GspD_α_ and GspD_β_ in the translocation process, with resolution ranging from 9 Å to 19 Å. In our results, GspD_α_ and GspD_β_ present entirely different membrane interaction patterns and ways of going across the peptidoglycan layer. We propose two distinct models for the membrane translocation of GspD_α_ and GspD_β_, providing a comprehensive perspective on the inner to outer membrane biogenesis of T2SS secretins.

## Introduction

Secretion systems, which are bacterial cell envelope-located protein complexes, are utilized by bacteria to produce virulence-related substrates that facilitate survival and pathogenicity^1^. Among secretion systems, the type II secretion system (T2SS) is broadly present and functional in *Proteobacteria* species, including non-pathogenic *Escherichia coli* (*E. coli*), Enterotoxigenic *E. coli* (ETEC), Enteropathogenic *E. coli* (EPEC), *Vibrio cholerae, Klebsiella pneumoniae*, and *Aeromonas hydrophila*^2^. T2SS substrates in non-pathogenic bacteria can facilitate nutrient absorption from the environment or symbiosis with plants or animals, whereas in pathogenic bacteria, T2SS substrates can aid in adhesion to hosts, intoxicate host cells, and suppress immunity in the host, causing various diseases^3^. With T2SS’s diverse substrate functions and close relevance to virulence and diseases, knowing its structure and working mechanism is necessary for understanding bacteria functions and developing new antimicrobial strategies.

The outer membrane component of T2SS is the secretin, constituting a large channel structure, connecting to the protein scaffold on the inner membrane, and controlling the last step of substrate transportation^2^. Phylogenetic analysis of T2SS secretins from *Proteobacteria* species demonstrated two types: the *Klebsiella*-type secretins found in *Klebsiella* and *Dickeya*; and *Vibrio*-type secretins found in *Vibrio*, ETEC, and EPEC^4^. Structures of the two secretin types both appear as cylindrical channels containing the N0-N3 domains, the secretin domain with a central gate region, and the S domain^5–10^. But they are different in the transmembrane region: the *Vibrio*-type secretins have about 20 additional amino acids between the α7 and α8 helices, forming a loop extending upward and inward to the central axis of the channel like a roof, constituting a cap gate, while *Klebsiella*-type secretins do not have the cap gate^5–10^. This directly causes differences in the height of the proposed transmembrane regions of the two secretin types, which may give them different ways of interacting with the membrane. T2SS secretins’ interactions with the membrane have not been clearly visualized either *in situ* or in purified systems, limiting our understanding of the substrate transportation mechanism and limiting options for drug inhibitor design. Furthermore, *in situ* structures of secretins are needed to understand how they assemble and function in living bacteria, in the context of the complete periplasmic/membrane environment. This environment cannot be accurately simulated *in vitro* and may have a significant impact on secretin behavior.

The translocation of T2SS secretins from the inner membrane to the outer membrane is an essential step required for T2SS assembly, however, it is still unclear how this translocation is regulated in the bacterial cell envelope. Based on biochemistry experiments, there exist two possible pathways: 1) the scaffolding proteins GspA and GspB bind and increase the pore size of peptidoglycan, and translocate the secretin to the outer membrane, as found in *Klebsiella*-type secretins ExeD and OutD^11–13^; 2) a small pilotin GspS could bind to the S domain of secretin, and the pilotin itself can translocate to the outer membrane through the Lol sorting pathway, as found in the *Vibrio*-type secretins^4,14,15^. Despite the current biochemical evidence, this translocation process has not been visualized in living cells. By doing *in situ* electron cryotomography (cryo-ET) studies on T2SS secretins, we capture different intermediate states during the outer membrane translocation process, which helps to explain the translocation mechanism and provide a clearer blueprint of the biogenesis process of T2SS secretins.

There are two T2SS secretins encoded in the *E. coli* genome: the *Klebsiella*-type GspD_α_ that can secret chitinase^16–19^, representing non-pathogenic functions; and the *Vibrio*-type GspD_β_^4,14^, which can secret toxins^20,21^, representing pathogenic functions. In this study, we investigate the *in situ* structures of GspD_α_ from the *E. coli* K12 strain and GspD_β_ from the ETEC H10407 strain as representatives of the *Klebsiella*-type secretins and *Vibrio*-type secretins, respectively. We report four *in situ* structural states of specific secretin/secretin-pilotin complex, determined within *E. coli* cells, using the cryo-ET subtomogram averaging. They are, respectively, GspD_α_ on the inner membrane, GspD_α_ on the outer membrane, GspD_β_-GspS complex on the outer membrane, and GspD_β_ on the inner membrane (Supplementary Table 1). Together, these structures show interactions of secretins with inner and outer membranes and provide new insights into the biogenesis process of the GspD secretin.

## Results

### Visualization of GspD_α_ on the bacterial inner membrane through cryo-ET

To obtain thin cells for improved contrast under cryo-ET, we employed a bacteria minicell system to provide thin cells which retain physiological activities^22^. We induced overexpression of GspD_α_ within *E. coli* BL21 (DE3) cells (Figure 1J). Cryo-ET was performed to image the minicells and 250 tilt series were collected. From the raw tilt images and the reconstructed tomograms, cell features including the intact bacterial cell envelope and membrane-bound GspD_α_ particles could be clearly recognized (Figure 1A-D). GspD_α_ multimer particles in different orientations with respect to the grid plane could be identified in the tomograms, and surprisingly, under these experimental conditions, the GspD_α_ particles are naturally located on the inner membrane of the bacterial envelope, with the non-transmembrane domains facing the periplasmic space (Figure 1D). As GspD_α_ has an approximately cylindrical shape, we identified particles in the top, side, and tilted views as in circles, a pair of curved lines attaching to the inner membrane, and parentheses object, respectively. The images of negative control cells further validated that these particle features found in the tomograms belong to our protein of interest (Supplementary Figure 1).

**Figure 1.**
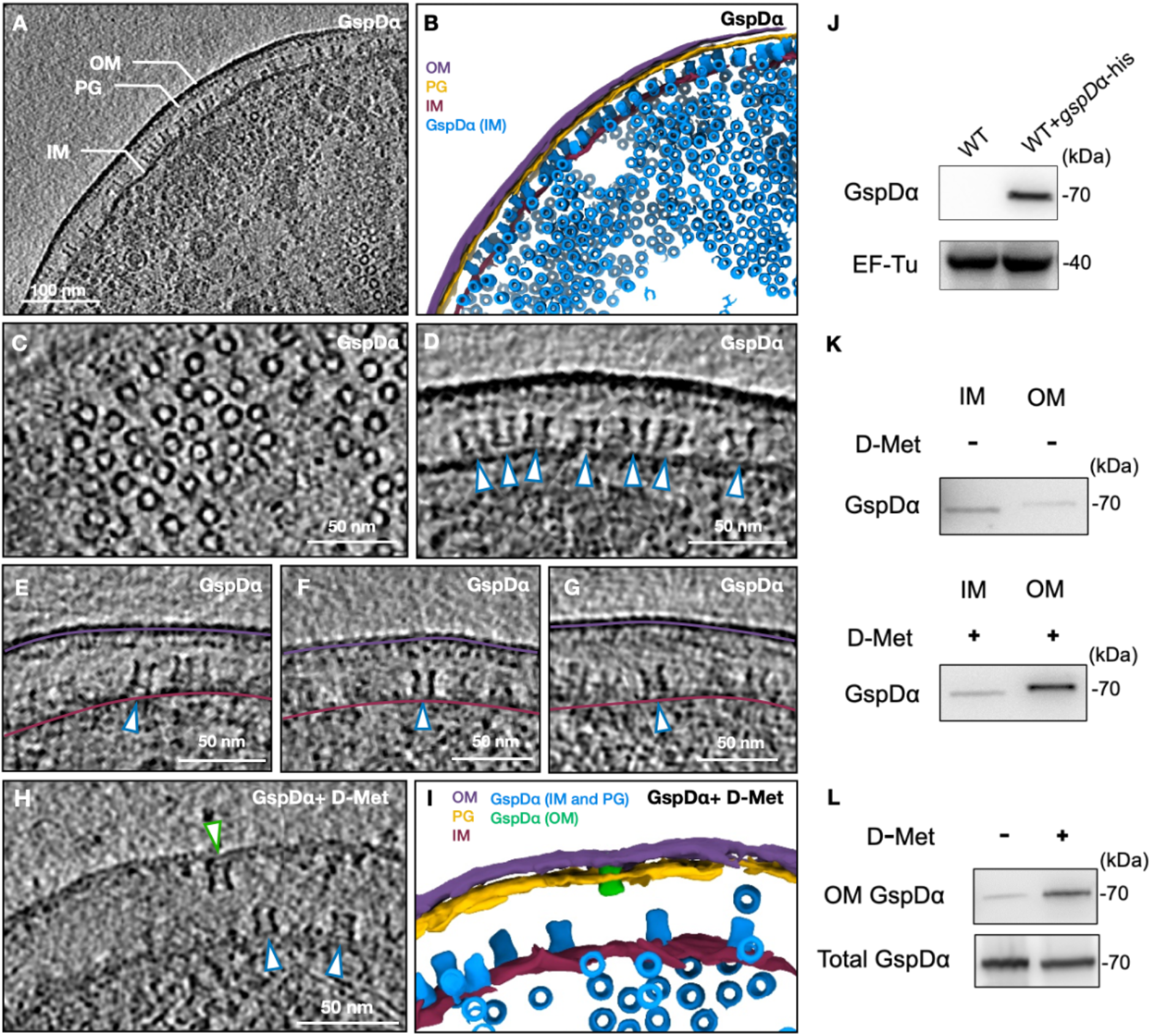
Visualization of GspD_α_ multimers within *E. coli* cells and determination of the overexpression and membrane location of GspD_α_. A, The tomogram z slice view of *E. coli* overexpressing GspD_α_. B, Segmentation of the tomogram shown in A. C, Zoom in tomogram z slice view of top view GspD_α_ particles. D, Zoom in tomogram z slice view of side view GspD_α_ particles (white arrowheads with blue outline). E, F, and G, Tomogram z slices showing tilted GspD_α_ on the inner membrane or GspD_α_ with only one side connecting the inner membrane (white arrowheads with blue outline). The outer and inner membranes are indicated by the purple line and the dark-red line, respectively. H, A tomogram z slice view of *E. coli* overexpressing GspD_α_ and with D-methionine added. I, Segmentation of the tomogram shown in H. Plasmid pETDuet-*gspDα*-his was transformed into BL21 (DE3) cells. Protein was induced by 0.5 mM IPTG at 20°C overnight. J, Immunoblotting results of GspD_α_ overexpression by using anti-His tag antibodies. BL21 (DE3) cells (WT) without plasmid transformation were used as the control. K, Inner and outer membrane fractions of the GspD_α_ overexpressing cells treated with/without D-methionine were separated by sucrose density gradient centrifugation. The membrane location of GspD_α_ was immunoblotted with anti-His tag antibodies. L, Outer membrane proteins of the GspD_α_ overexpressing cells treated with/without D-methionine were extracted using a bacterial membrane protein extraction kit. The total GspD_α_ and outer membrane GspD_α_ were examined by western blot analysis using anti-His tag antibodies. IM, inner membrane; OM, outer membrane; PG, peptidoglycan; D-Met, D-methionine; WT, wild type.

### *In situ* subtomogram averaging structure of GspD_α_ at subnanometer resolution

For subtomogram averaging, ∼32,000 particles were manually picked, and data processing was carried out using EMAN2^23^. In previous *in vitro* structures of secretins, different symmetries have been reported, including C12 and C15^5,24,25^. To confirm the symmetry of our particles, we developed an algorithm to measure the radius of individual particles and generated a histogram to look for potential multiple radii (Supplementary Figure 2). The histogram shows only one peak with a diameter consistent with the C15 structure of GspD_α_ (PDB: 5WQ7), strongly implying that there is only one diameter of GspD_α_ particles *in situ* on the inner membrane, corresponding to its C15 structure. One challenge in subtomogram alignment of the secretin is that, due to the lack of low-resolution features along the symmetry axis, at the initial stage of orientation determination, it is difficult to distinguish top-view particles from bottom-view ones without information about the membrane position. Therefore, at the beginning of the subtomogram refinement, many top-view particles were misaligned and positioned upside down, limiting the final resolution, and causing the initial reconstructed map to exhibit some “D” symmetric features in addition to the 15-fold symmetry. To resolve this ambiguity, we developed an algorithm in the EMAN2 tomography package, which considers the geometric location of the cell membrane as an additional factor in particle orientation determination (see Methods). The use of this methodology improved both visible features in the map as well as the measured resolution (Supplementary Figure 3B-C).

The final C15 symmetrized subtomogram average achieved a measured resolution of 9 Å (Supplementary Figure 4A), demonstrated by some visibility of alpha helices in the map (Supplementary Figure 5B, Supplementary Movie 1). The inner membrane bilayer is clearly resolved and visible at the bottom of this density map. Above the inner membrane bilayer is the spool-shaped protein (cylinder with a larger diameter at both ends) with a sealed membrane adjacent surface and an open membrane distal surface. The N0, N1, N2, and N3 domains can be clearly recognized in the density map (Supplementary Figure 5B). In N1 and N2 domains, the two layers of alpha-helices are just resolved, consistent with the measured resolution of the map. With the position of the N domains identified, the transmembrane region of GspD_α_ can be approximately modeled using the length of the secretin domain. The tip of GspD_α_ (α7, α8, α9, α10, β11, β14, and the membrane-buried region of β10 and β15^5^) transits through one leaflet of the lipid bilayer but does not penetrate through or generate an opening on the membrane (Supplementary Figure 5B). The sealed end of the cylinder corresponds to the gate region of the secretin. The connecting density between the protein and the inner membrane appears to have very low occupancy based on the low isosurface threshold required to visualize it. Interpreting this connection thus required some additional effort.

### GspD_α_ is partially connected to and flexible on the bacterial inner membrane

As both the membrane and GspD_α_ particles are clearly resolved in the raw tomogram, we could directly observe that some particles were slightly tilted with respect to the membrane with a connection of varying strength (Figure 1E-G). To further confirm the membrane connection to GspD_α_, using the C15 density map and the corresponding orientation information of each particle, we generated an asymmetric structure by relaxing the symmetry (Supplementary Figure 3E-F) (see Methods), producing a structure with a reduced resolution of 15 Å (Supplementary Figure 5D). In this density map, the inner membrane leaflets are still resolved, as expected, and GspD_α_ constitutes the middle-contracted cylindrical density. The membrane connecting density is now clearly visible with apparent full occupancy, but only on one side of the cylinder. The membrane connection covers a roughly 100-degree arc, depending on the threshold level, so when bound to the inner membrane, GspD_α_ is only partially connected. When C15 symmetry is applied along the cylindrical axis, this partial connection is averaged out, yielding the lower apparent occupancy in the original symmetrized structure (Supplementary Figure 5B).

In addition, we performed a focused refinement on the protein region, excluding the membrane, and compared the particle orientation with that of the integrated refinement result (Supplementary Figure 3G) (see Methods). The orientation difference between the two refinements can then be determined for each particle. By classifying these differences, we recovered a trajectory for GspD_α_ on the inner membrane, where the multimer swings around the membrane contact site (Supplementary Movie 2). We show the two endpoints’ conformations for this trajectory (Figure 2B-C): in conformation A, the symmetry axis of GspD_α_ is perpendicular to the membrane; and in conformation B, the symmetry axis of GspD_α_ is tilted 2.86 degrees compared to conformation A.

**Figure 2.**
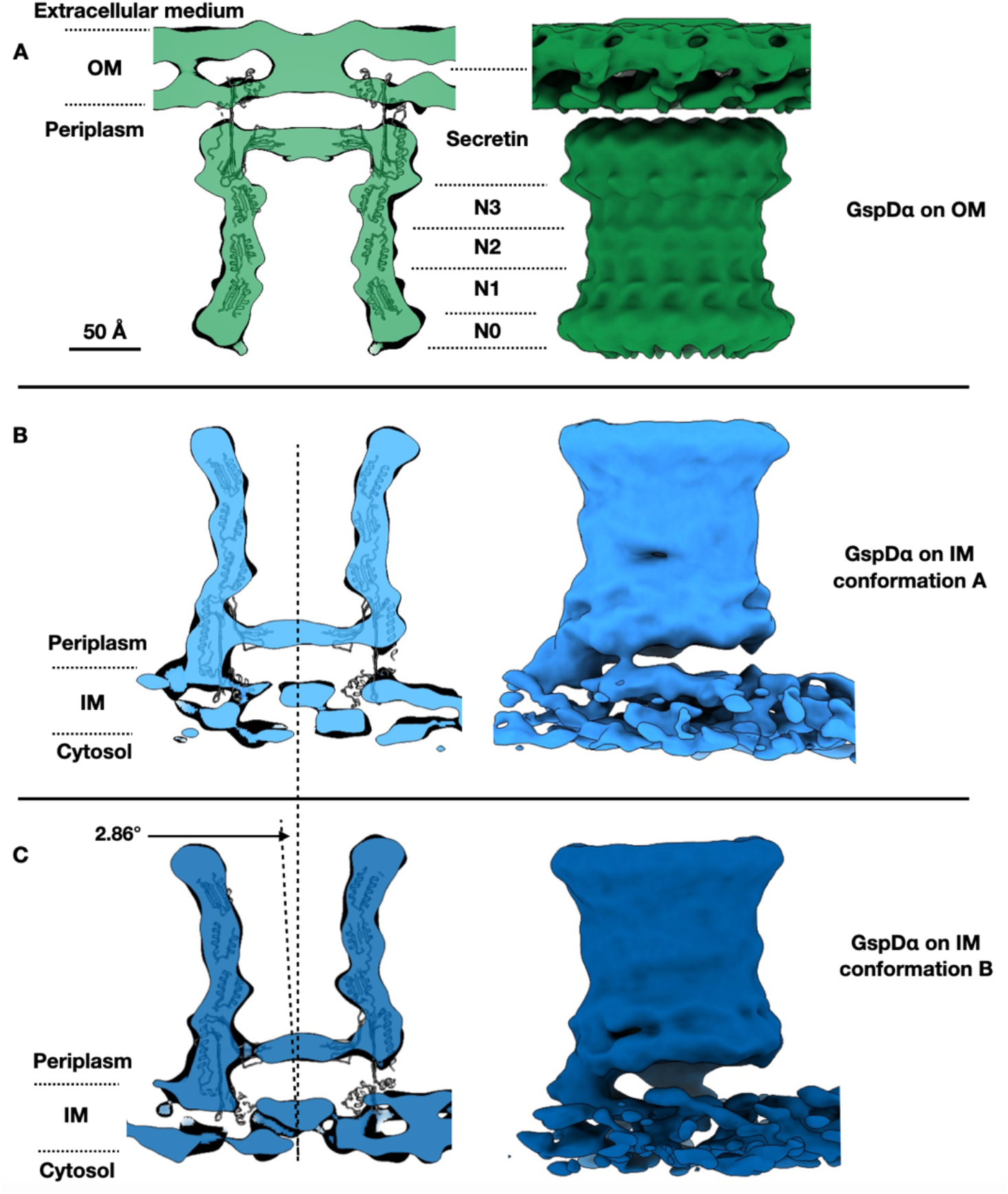
*In situ* structures of the GspD_α_ secretin on the outer and inner membranes. A, The *in situ* structure of GspD_α_ on the outer membrane, showing the central slice view and side view. B and C, The *in situ* structures of GspD_α_ conformation A and conformation B on the inner membrane respectively, showing the central slice view and side view. Density maps are fitted by the GspD_α_ *in vitro* structure (PDB: 5WQ7). OM, outer membrane; IM, inner membrane.

In summary, these results indicate that the GspD_α_ multimer is not integrated into the bacterial inner membrane. Instead, it only forms a loose connection with significant mobility, facilitating its further transportation to the outer membrane (see Discussion).

### GspD_α_ is inserted into the bacterial outer membrane when raising *E. coli* with D-methionine added

We hypothesize that the GspD_α_ multimer is on the inner membrane because of the existence of the peptidoglycan, which is a meshwork whose pore size is too small for the GspD_α_ multimer to pass through and translocate to the outer membrane. We attempted to reduce peptidoglycan crosslinking to see if this would facilitate GspD_α_ outer membrane targeting. In previous studies, it was shown in *Aeromonas hydrophila* that decreasing the cross-linking of the peptidoglycan by glycine could localize the secretin ExeD to the outer membrane^11,12^. Biochemical evidence also showed that the growth of *E. coli* in media containing a high concentration of D-methionine decreases the cross-linking of the *E. coli* peptidoglycan^26^. Therefore, we raised *E. coli* minicells in LB medium containing 40 mM D-methionine, induced expression of GspD_α_, and found that this increased the proportion of GspD_α_ on the outer membrane compared to the control (Figure 1K, L). Within the tomograms, we observed GspD_α_ particles located on the outer membrane as well as the inner membrane (Figure 1H-I). For subtomogram averaging, we only selected particles on the outer membrane. There were 309 particles, leading to a final refinement with C15 symmetry at 16 Å resolution (Supplementary Figure 4B). Within the density map, the outer membrane bilayer is again resolved with an adjacent spool-shaped density. The membrane adjacent surface of the cylinder is sealed, indicating the gate region. The N0, N1, N2, and N3 domains could be recognized from the membrane distal to the membrane adjacent side of the cylinder (Figure 2A).

To further investigate the protein-membrane interactions, we relaxed the C15 symmetry as previously for GspD_α_ on the inner membrane. The resulting density map, seen from the cross-section at the membrane connection, shows GspD_α_ connecting to the membrane with an even distribution of occupancy on the contour of the cylinder periphery (Supplementary Figure 5C), instead of the one-side connection observed on the inner membrane (Supplementary Figure 5D). To examine the symmetry of GspD_α_ on the outer membrane, we did a refinement imposing only C5 symmetry, and the density map retains clear C15 features (Supplementary Figure 5E), confirming the symmetry of GspD_α_ *in situ*. For comparison, we did C5 refinement on the GspD_α_ on the inner membrane dataset, and the resulting density map did not manifest clear C15 features (Supplementary Figure 5F). Together, these results show that enlarging the pore size of peptidoglycan by adding D-methionine when raising *E. coli* permits GspD_α_ to translocate to the outer membrane, where GspD_α_ adopts a more consistent conformation and forms an evenly-distributed connection with the membrane.

### GspD_β_ on the bacterial outer membrane co-exists with GspS

GspD_β_, as a homolog of GspD_α_, has a similar structure except for an additional cap gate on the membrane adjacent side^5,8^, which may generate different transmembrane regions and membrane interactions. Also, GspD_β_ has a different outer membrane targeting mechanism. Instead of using scaffolding proteins or enlarging the peptidoglycan pore size, a small protein GspS, named pilotin, could bind to the S domain of GspD_β_ and translocate GspD_β_ to the outer membrane^4,14^. Therefore, to visualize GspD_β_ particles on the outer membrane and verify whether GspS forms a complex with GspD_β_ there, we performed two experiments. One is that the GspD_β_ and GspS expression are both induced in wild-type *E. coli* minicells, the other one is that only GspD_β_ expression is induced in wild-type *E. coli* minicells; and the protein expression was verified (Figure 3H).

**Figure 3.**
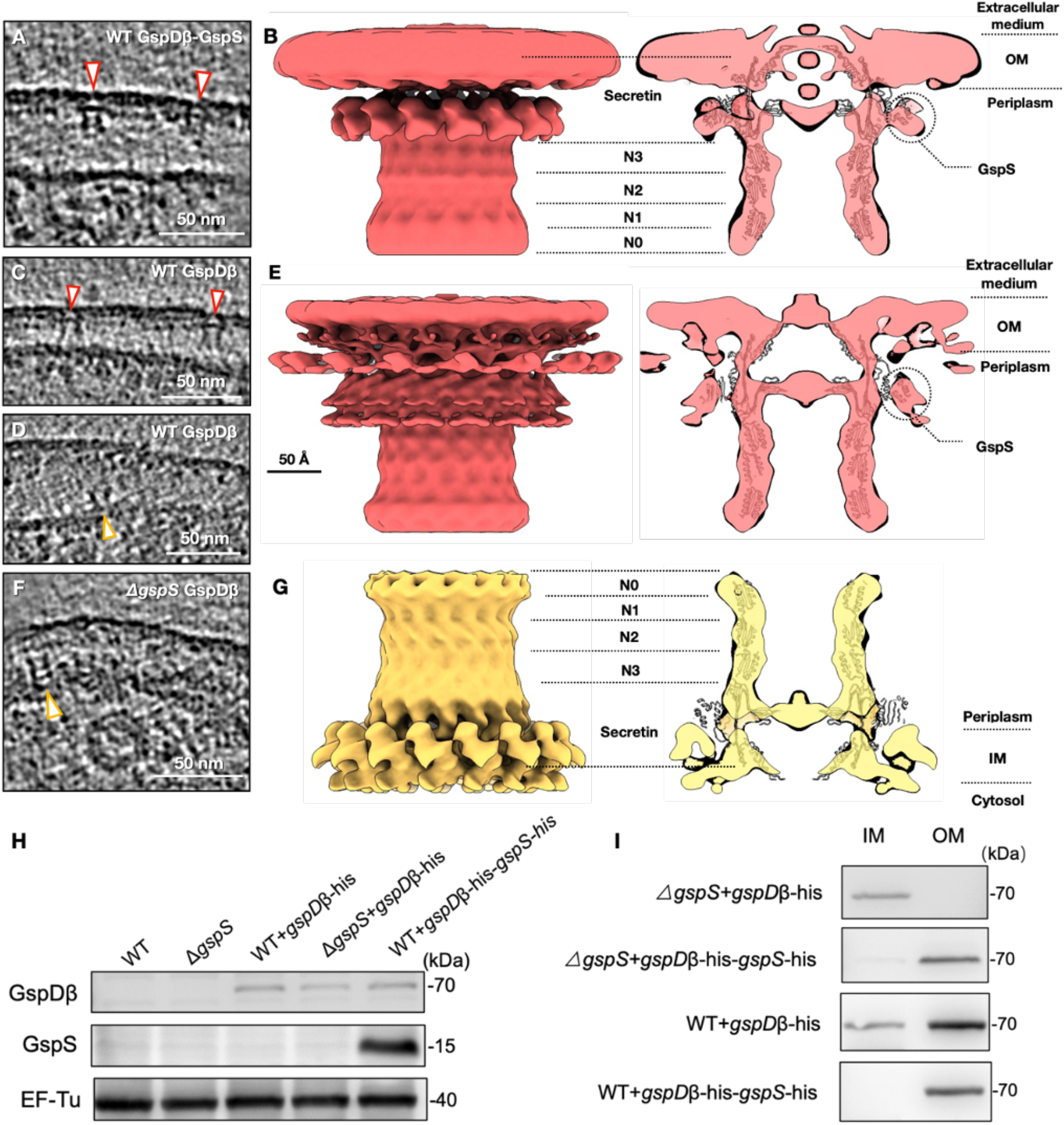
*In situ* structures of the GspD_β_ secretin and determination of the overexpression and membrane location of GspD_β_. A, The tomogram z slice view of *E. coli* overexpressing GspD_β_ and GspS. B, The *in situ* structure of the GspD_β_-GspS complex on the outer membrane when both GspD_β_ and GspS are overexpressed, showing the side view and central slice view. C and D, Tomogram z slice views of *E. coli* overexpressing only GspD_β_. E, The *in situ* structure of the GspD_β_-GspS complex on the outer membrane when only GspD_β_ is overexpressed, showing the side view and central slice view. F, The tomogram z slice view of *∆gspS E. coli* overexpressing GspD_β_. G, The *in situ* structure of the GspD_β_ multimer on the inner membrane, showing the side view and central slice view. Particles on the outer membrane and the inner membrane are indicated by white arrowheads with a red outline and white arrowheads with a yellow outline, respectively. The density maps are fitted with the *in vitro* structure of the GspD_β_-GspS complex (PDB: 5ZDH). H, Immunoblotting results of GspD_β_ and GspS overexpression by using anti-his antibodies. BL21 (DE3) cells (WT) and BW25113-Δ*gspS* cells (Δ*gspS*) without plasmid transformation were used as the control. I, Detection of the membrane location of GspD_β_ through sucrose density gradient centrifugation followed by western blot analysis using anti-His tag antibodies (the bacteria strains are marked on the left, and all the bands correspond to GspD_β_). IM, inner membrane; OM, outer membrane; WT, wild type.

In the reconstructed tomograms of GspD_β_-GspS expressed cells, particles are all found on the outer membrane (Figure 3A), verified by a membrane separation experiment (Figure 3I). We boxed 514 subtomogram particles, and the refinement with C15 symmetry achieved a 19 Å resolution (Supplementary Figure 4C). To verify the symmetry, we did a refinement with C5 symmetry applied, and the resulting density map still shows C15 features (Supplementary Figure 6A), confirming that GspD_β_ *in situ* has C15 symmetry. In the density map (Figure 3B), the outer membrane density is visible as the top layer. Below the outer membrane is the spool-shaped density, similar to that of GspD_α_. The membrane adjacent surface of the cylinder density is sealed, indicating the gate region. Notably, extra density appearing as 15 lumps can be seen connecting to the membrane adjacent periphery of the cylinder. These extra densities likely belong to GspS, indicating that GspD_β_ on the outer membrane exists in a complex with GspS. The *in vitro* structure of the GspD_β_-GspS complex (PDB: 5ZDH) could be well fitted into the density map, and according to the position of the membrane density, the transmembrane region could be located at α7, α8, α9, α10, β11, β12, β13, β14, and the membrane buried region of β10 and β15^5,8^. In contrast to GspD_α_ which only transits through one leaflet of the membrane, GspD_β_ transit through two leaflets of the lipid bilayer, with the additional cap gate contacting the outer leaflet of the outer membrane.

We expect that in cells where only GspD_β_ is expressed, particles will be on the outer membrane since there should be the endogenous expression of GspS in the *E. coli* BL21(DE3) strain. Observed from the tomograms, most particles are on the outer membrane (Figure 3C), but surprisingly, a few particle side views are seen on the inner membrane (Figure 3D), which was verified with a membrane separation experiment (Figure 3I). Within this dataset, 910 particles were picked, including a mixture of outer and inner membrane-located particles. After one round of refinement, we performed a multi-reference classification with two references rotated 180 degrees with respect to each other, which produced two populations of particles with a ratio of 723 outer membrane particles to 187 inner membrane particles. We used the 723 particles to do the refinement with C15 symmetry and achieved a 16 Å resolution structure (Figure 3E and Supplementary Figure 4D). Refinement with C5 symmetry was also performed, and the density map showed C15 features (Supplementary Figure 6B). This structure basically resembles that of the GspD_β_-GspS expressed dataset (Figure 3B), but notably, the GspS density is still seen outside the GspD_β_ channel, appearing as lumps connecting to the cylinder surface, although we did not induce overexpression of GspS.

Together, these results indicate that, when GspS is overexpressed together with GspD_β_, they form a complex on the outer membrane. Without exogenous GspS expression, when GspD_β_ is expressed, the *E. coli* endogenous GspS is sufficient to locate GspD_β_ to the outer membrane, where GspS stays attached to GspD_β_.

### GspD_β_ multimer is visualized on the bacterial inner membrane when *gspS* is knocked out

As biochemistry results have shown that GspD_β_ could form multimers on the inner membrane when *gspS* is knocked out^4,14^, to visualize GspD_β_ on the inner membrane, we induced expression of GspD_β_ within Δ*gspS E. coli* cells (Figure 3H). Within the reconstructed tomograms, particles are all visualized on the inner membrane, with the non-transmembrane domains located in the periplasm (Figure 3F, I). There are 370 particles, and the refinement with C15 symmetry achieved a 14 Å resolution structure (Supplementary Figure 4E), rendering GspD_β_ on the inner membrane (Figure 3G). The membrane density could be located at the bottom, and the cylinder density appears above, inserted into the membrane. The sealing at the lower end of the cylinder corresponds to the gate region. The lower end of GspD_β_ transits through two leaflets of the inner membrane, and there is no GspS density seen outside the cylinder density. These results indicate that, when *gspS* is knocked out, GspD_β_ could self-assemble into a multimer that has stable conformation on the inner membrane.

## Discussion

In this study, we show the *in situ* structures of the GspD_α_ secretin from bacterial T2SS and their interactions with the inner and outer membranes. We show that: 1) GspD_α_ could self-assemble into its multimeric form on the inner membrane without overexpression of any other protein (Figure 1D); 2) GspD_α_ multimer forms flexible connections with the inner membrane, and does not generate an opening on the inner membrane (Figure 2B-C); 3) the undisturbed normal peptidoglycan pore size does not allow the passage of a GspD_α_ multimer^27^; 4) dissociative GspD_α_ multimers exist in the periplasm of *E. coli* expressing GspD_α_, but only between the peptidoglycan and the inner membrane (Supplementary Figure 7A-C); 5) D-methionine could reduce peptidoglycan crosslinking^26^, and adding D-methionine when raising *E. coli* could localize GspD_α_ to the outer membrane (Figure 1H); 6) dissociative GspD_α_ multimers exist in the periplasm of *E. coli* raised with D-methionine and expressing GspD_α_ (Supplementary Figure 7D); 7) GspD_α_ on the outer membrane exists in a more stable conformation, showing distinguishable symmetry, and forming evenly-distributed connections with the outer membrane (Supplementary Figure 5C and 5E). Altogether, this evidence supports a translocation model in which GspD_α_ first forms multimers on the inner membrane, where it stays in an intermediate state, unstable and swinging, which could favor its transportation to the outer membrane. When the peptidoglycan pore is enlarged by D-methionine, the multimer translocates to the outer membrane without disassembly. On the outer membrane, it exists in a consistent conformation, firmly attached to the membrane, which potentially facilitates substrate transportation (Figure 4A).

**Figure 4.**
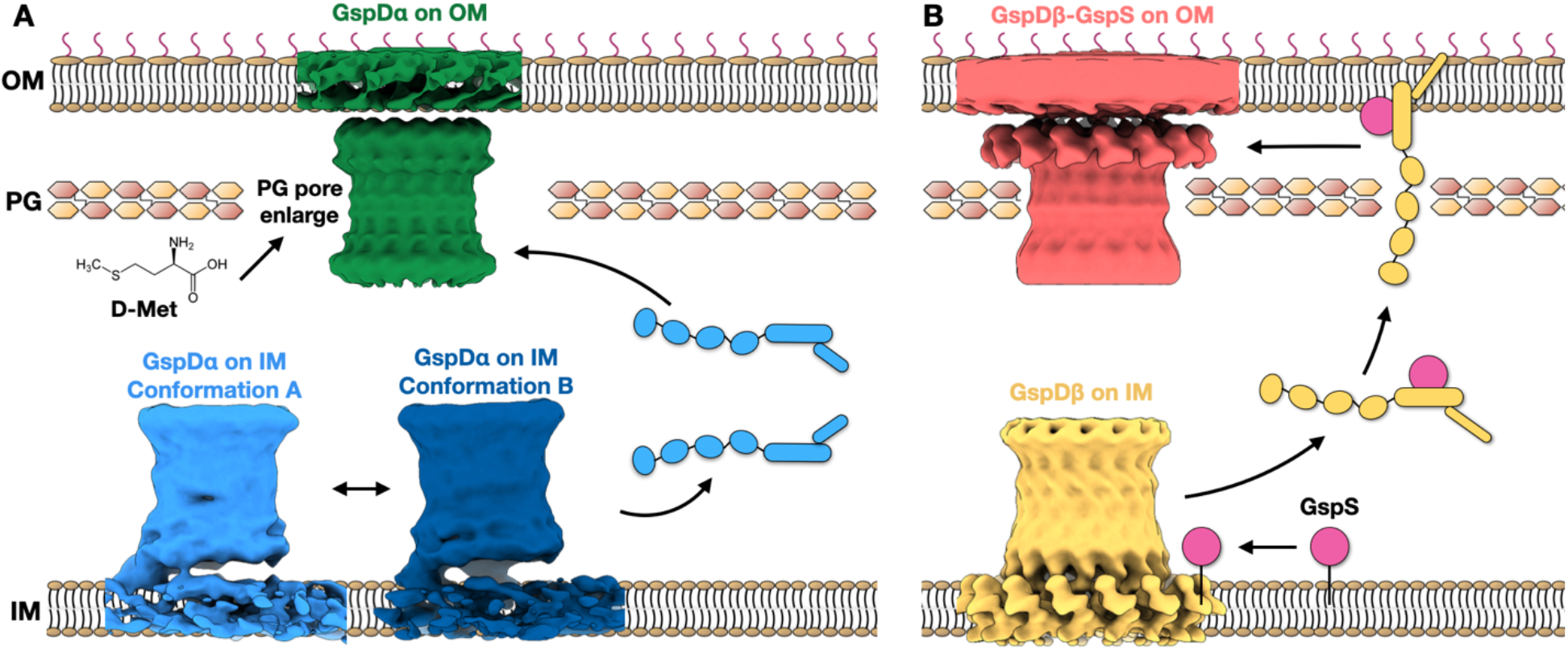
Proposed models of the outer membrane translocation process of the GspD_α_ secretin (A) and the GspD_β_ secretin (B). A, Firstly, GspD_α_ forms multimers on the inner membrane, and adopts an unstable and swinging conformation, changing between conformation A (light blue) and conformation B (dark blue). The multimer could dissociate from the inner membrane and enter the periplasm (light blue cartoon model), but it could not pass through the peptidoglycan and insert into the outer membrane. When the peptidoglycan pore size is enlarged by D-methionine, the multimer translocates to the outer membrane, where it exists in a more stable conformation (green). B, GspD_β_ first forms multimers on the inner membrane; then the pilotin GspS approaches and binds the GspD_β_ monomer, transporting it to the outer membrane. On the outer membrane, GspD_β_ assembles into a multimer again and GspS stays associated, forming a GspD_β_-GspS multimer complex. IM, inner membrane; PG, peptidoglycan; OM, outer membrane; D-Met, D-methionine.

We observed GspD_β_ multimer structures on both the bacterial inner and outer membranes. We show that: 1) when *gspS* is knocked out in *E. coli*, GspD_β_ forms multimers on the inner membrane (Figure 3F-G)^4^; 2) when *gspS* is not knocked out or overexpressed in *E. coli*, a small number of GspD_β_ multimers are still found on the inner membrane (Figure 3D); 3) the undisturbed normal peptidoglycan pore size does not allow the passage of GspD_β_ multimers^27^; 4) when GspS exists, either from vector overexpression or endogenous genome expression, GspD_β_ multimers are found on the outer membrane (Figure 3A and 3C), even if the peptidoglycan is undisturbed; 5) GspS binds to GspD_β_ monomer in a 1:1 ratio^8^; 6) GspS itself is a lipoprotein that could translocate from the inner to the outer membrane through Lol pathway^14^; 7) we do not observe disassociated GspD_β_ multimers in the periplasm; 8) GspD_β_ multimers on the outer membrane exist in a complex with GspS, that is, GspS does not dissociate after transportation of GspD_β_ to the outer membrane (Figure 3B and 3E). Based on what we observed, we would propose a translocation model that, first GspD_β_ self-assembles into multimers on the inner membrane, then GspS on the inner membrane binds to GspD_β_, which dissociates into monomers. GspS then transports GspD_β_ to the outer membrane, where GspD_β_ assembles into multimers again with GspS associated, forming a GspD_β_-GspS multimer complex (Figure 4B). Because of the limitations of cryoET observations, we did not directly visualize the process of GspD_β_ multimers on the inner membrane dissociating into monomers and entering the periplasm with GspS bound. Further investigations are needed to provide evidence for this process.

In our structures, the transmembrane region of GspD_α_ is only buried inside one leaflet of the membrane, instead of penetrating the membrane. This transmembrane pattern is unexpected and would be biophysically unreasonable but can be clearly observed from our cryo-ET structures. Further investigations are necessary in order to explain why this membrane interaction could exist. Comparing the inner and outer membrane-located structures of GspD_α_ and GspD_β_, we observe that GspD_α_ only transits one leaflet of the lipid bilayer, but GspD_β_ transits through both leaflets (Supplementary Figure 8). This difference could give GspD_β_ higher efficiency when transporting substrates. On the *E. coli* genome, there are two T2SS gene operons, T2SS_α_ and T2SS_β_. During evolution, T2SS_β_ may be generated by gene duplication of the T2SS_α_ operon. However, T2SS_α_ is endogenously silenced, while T2SS_β_ is actively expressed and functional^14^. Our result could provide a possible explanation, that is, the possibly higher substrate transport efficiency gives GspD_β_ an advantage during evolution, making it the preferred and expressed T2SS in many bacteria.

In summary, we determined four *in situ* structures of the T2SS secretin: both the inner and outer membrane structures of GspD_α_ and GspD_β_. GspD_α_ exists in an unstable form on the inner membrane, but forms firm and stable connections with the outer membrane. The GspD_β_ multimer exists on the inner membrane in wild-type *E. coli*, and GspD_β_ on the outer membrane exists together with GspS. Taken together, these results add to the knowledge of the membrane interactions of the T2SS secretins and their outer membrane targeting process, providing a comprehensive model for the secretin biogenesis process of *Proteobacteria*.

## Supplementary tables

**Supplementary Table 1.** Statistics of *in situ* structures achieved in this study. IM, inner membrane; OM, outer membrane.

| Structure | GspD<sub>α</sub> on IM | GspD<sub>α</sub> on OM | GspD<sub>β</sub>-GspS on OM | GspD<sub>β</sub>-GspS on OM | GspD<sub>β</sub> on IM |
| --- | --- | --- | --- | --- | --- |
| Overexpressed protein(s) | GspD<sub>α</sub> | GspD<sub>α</sub> | GspD<sub>β</sub> and GspS | GspD<sub>β</sub> | GspD<sub>β</sub> |
| Microscope | Titan Krios | Titan Krios | Glacios | Glacios | Glacios |
| Tilt scheme | Bidirectional | Bidirectional | Dose symmetry | Dose symmetry | Dose symmetry |
| Angstrom per pixel | 1.76 | 1.76 | 1.52 | 1.52 | 1.52 |
| Tilt series number | 250 | 91 | 63 | 191 | 24 |
| Particle number | 32,915 | 309 | 514 | 723 | 370 |
| Resolution (C15) | 9 Å | 16 Å | 19 Å | 16 Å | 14 Å |

## Supplementary figures

**Supplementary Figure 1.**
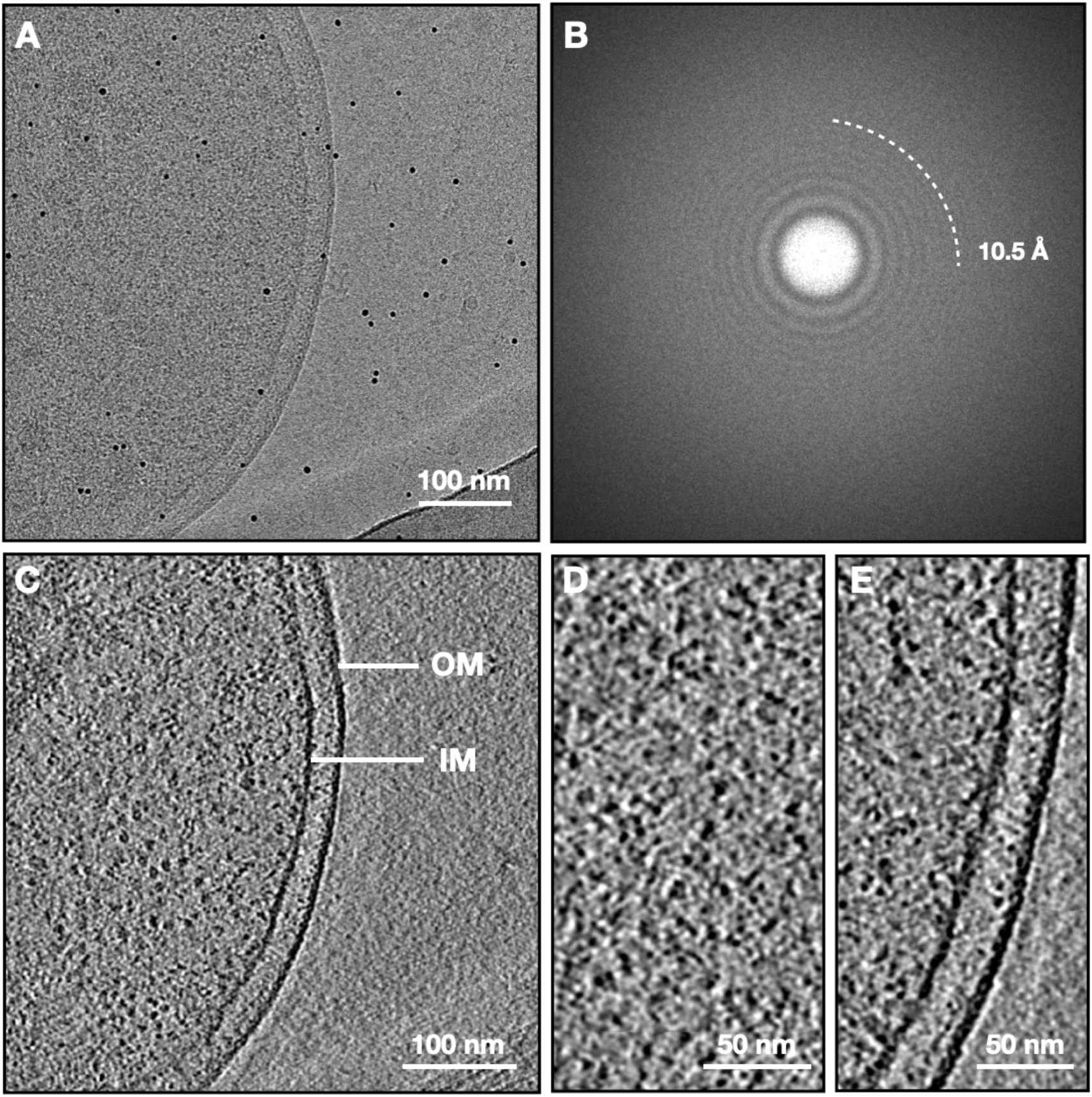
The cryo-ET tomogram of control *E. coli* minicells not overexpressing any protein. A, A tilt image at 0 degrees. B, The Fourier transform of A, showing the resolving ability. C, The reconstructed tomogram of an *E. coli* minicell not expressing any protein. D, Zoomed in tomogram z slice view of an area inside the cell body. E, Zoomed in tomogram z slice view of the cell envelope region. OM, outer membrane; IM, inner membrane.

**Supplementary Figure 2.**
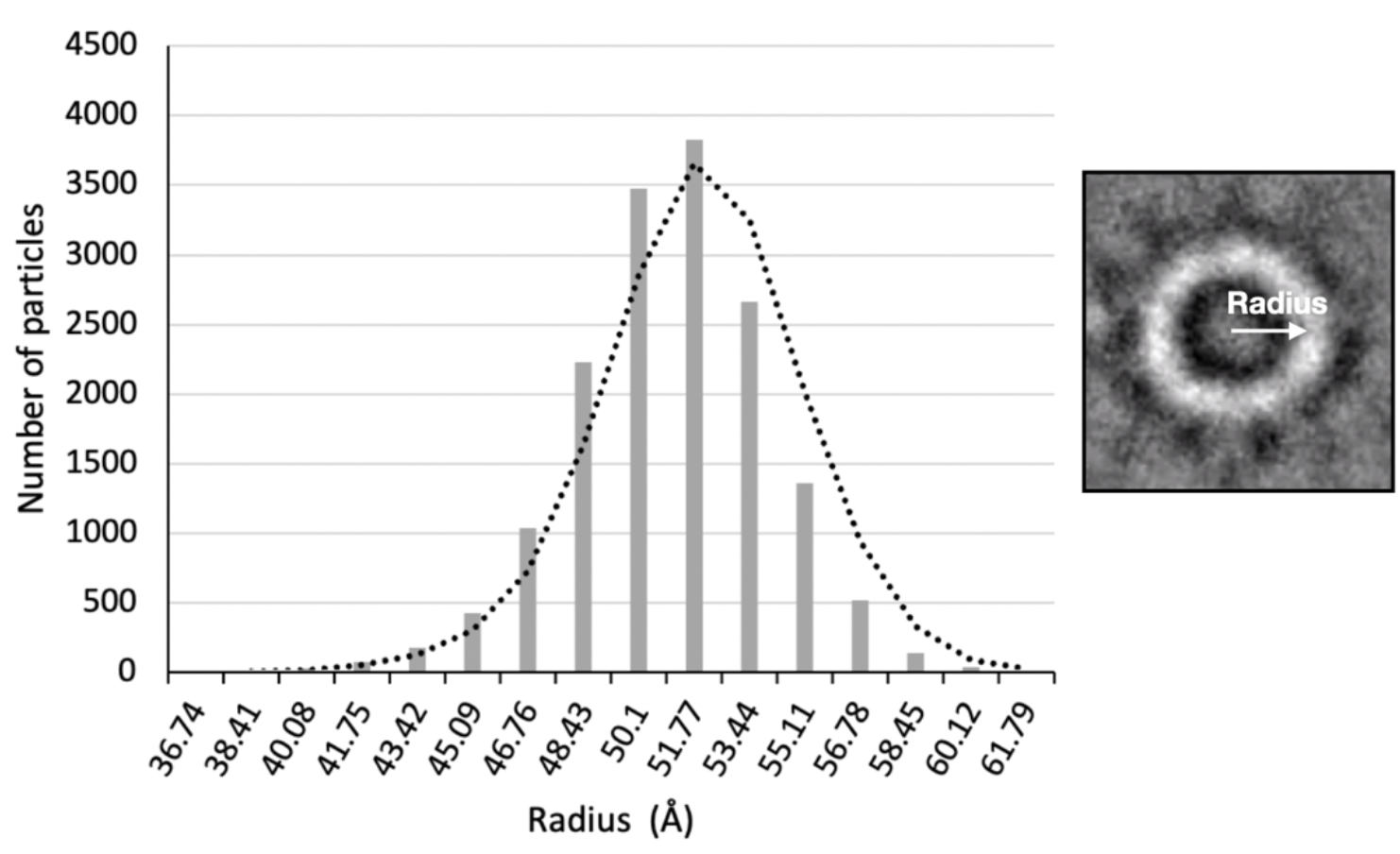
The histogram distribution of the radius of GspD_α_ particles *in situ*. The trend line is the moving average of the histogram data. A particle top view 2D image is shown on the right to illustrate the way of measuring the radius.

**Supplementary Figure 3.**
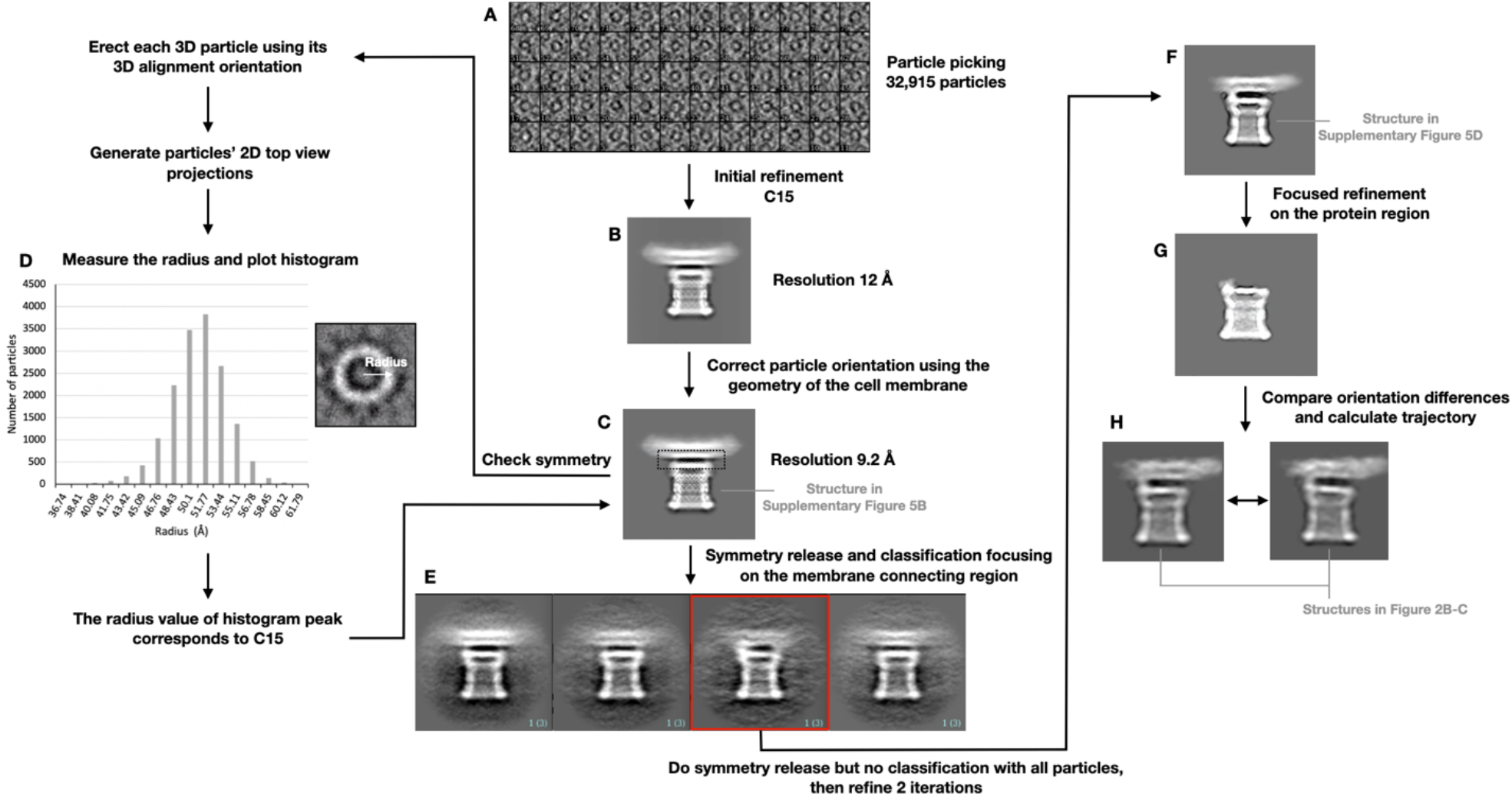
The data processing workflow for the *E. coli* overexpressing GspD_α_ dataset. After the manual boxing of particles (A), an initial refinement was done with C15 symmetry (B). Then the cell membrane feature was used to correct the particle orientations (see Methods). A local refinement was performed with the corrected orientation as input, achieving the 9.2 Å resolution structure (C). To verify the symmetry here, we measured and plotted the particle radius (D). The symmetry release and classification were focused on the membrane connecting region marked by the dashed line box in C, which resulted in four classes (E), including a class that shows an obvious uneven connection to the membrane (red line box in E). We then did a symmetry release with the whole dataset, followed by 2 iterations of refinement, producing the C1 structure (F). To demonstrate the particle movement, we performed a focused refinement with a mask only covering the protein density (G) and compared the orientation difference to that from the integral refinement. Conformations within this movement could be presented by calculating the trajectory and interval class averages on the trajectory (H).

**Supplementary Figure 4.**
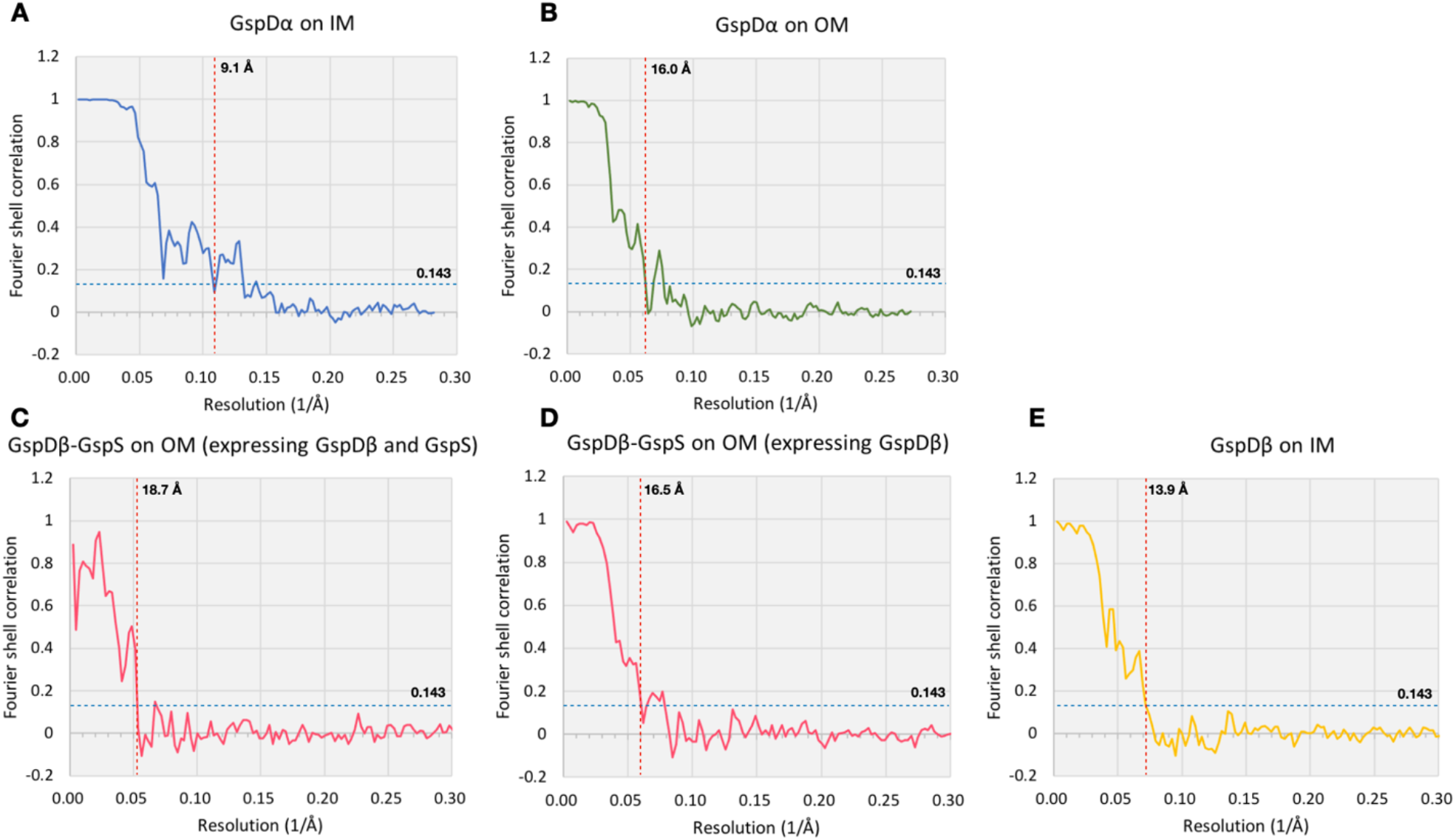
The FSC curves showing the resolution of *in situ* structures achieved in this work.

**Supplementary Figure 5.**
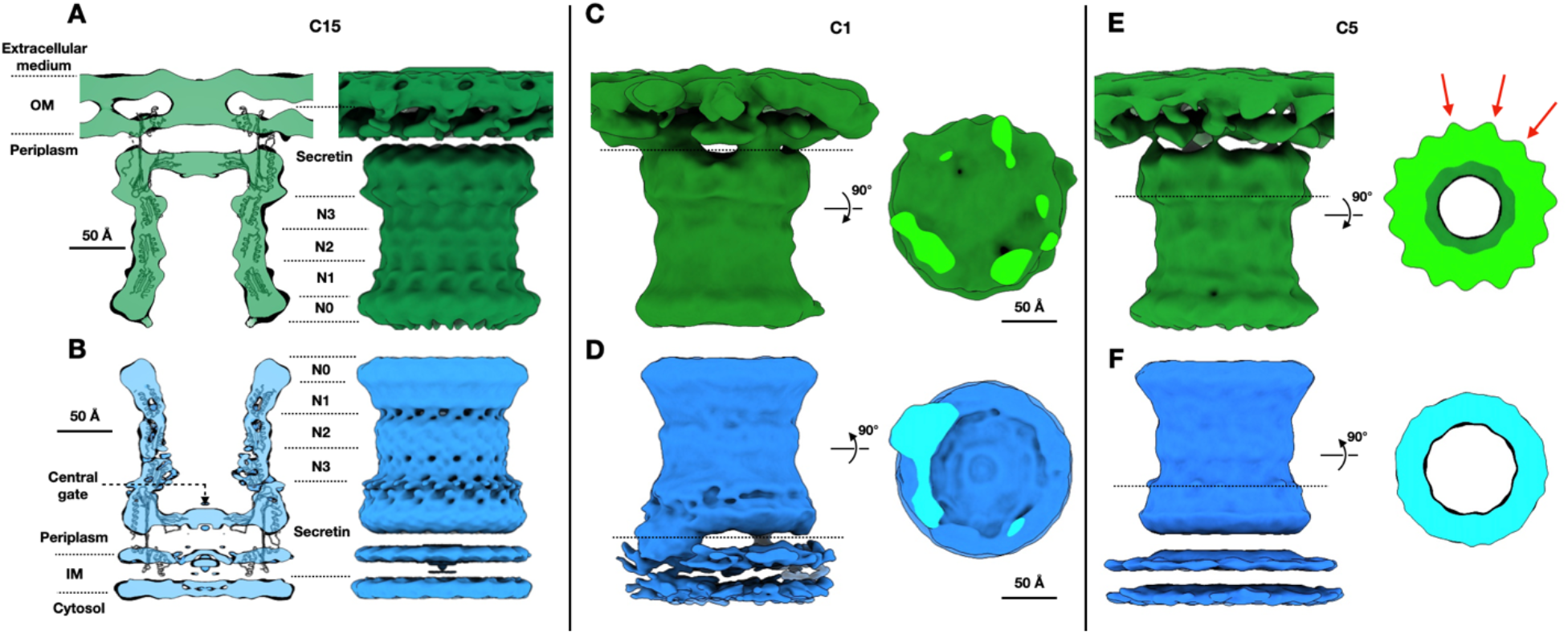
Comparison of GspD_α_ *in situ* structures on the inner and outer membranes. A, The *in situ* structure of GspD_α_ on the outer membrane, applied with C15 symmetry, fitted with GspD_α_ *in vitro* structure (PDB: 5WQ7). B, The *in situ* structure of GspD_α_ on the inner membrane, applied with C15 symmetry, fitted with GspD_α_ *in vitro* structure (PDB: 5WQ7). C, The *in situ* structure of GspD_α_ on the outer membrane, applied with C1 symmetry. D, The *in situ* structure of GspD_α_ on the inner membrane, applied with C1 symmetry. E, The *in situ* structure of GspD_α_ on the outer membrane, applied with C5 symmetry. The three units within one C5 symmetry unit are indicated by red arrows. F, The *in situ* structure of GspD_α_ on the inner membrane, applied with C5 symmetry. IM, inner membrane; OM, outer membrane.

**Supplementary Figure 6.**
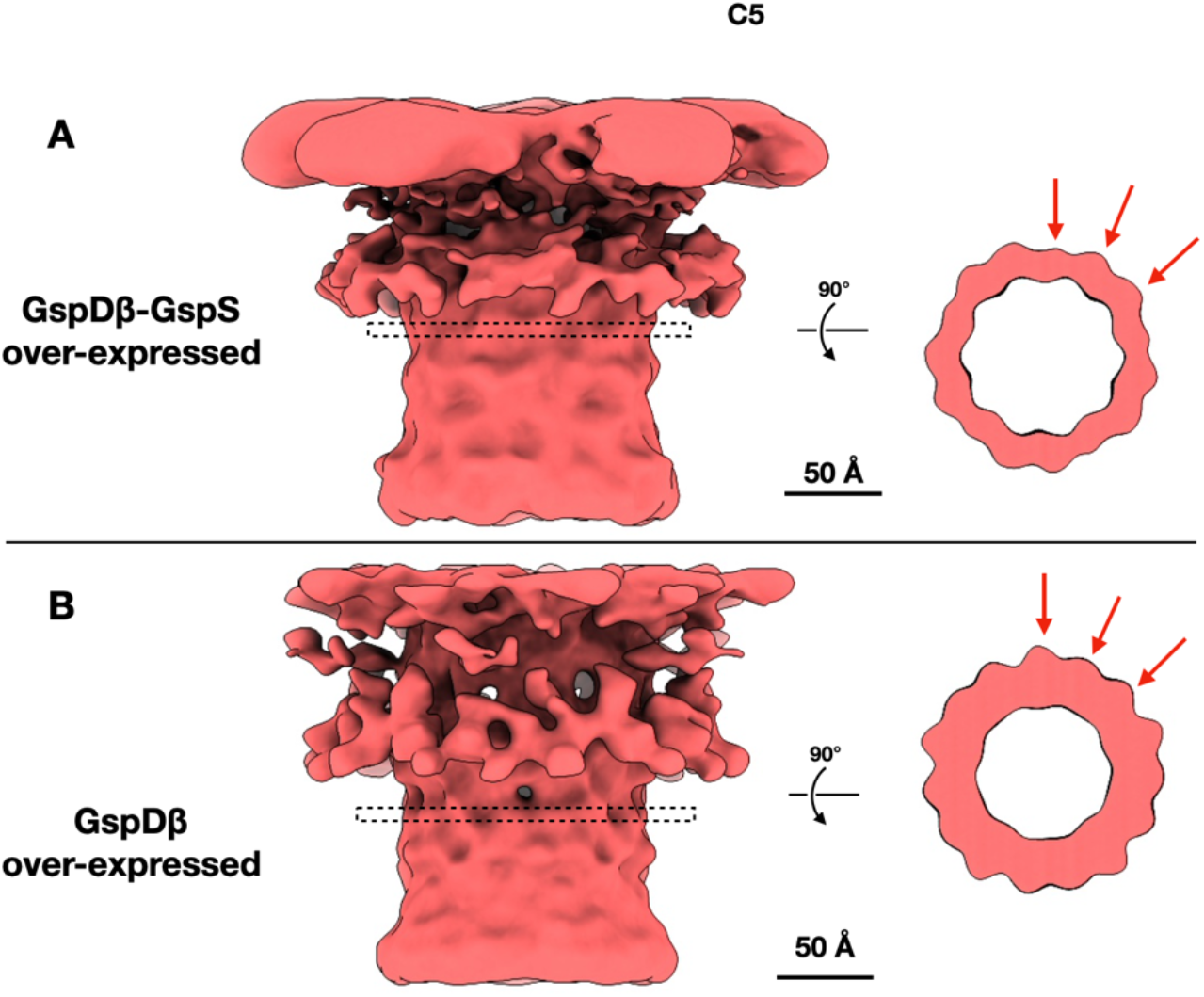
The *in situ* structures of the GspD_β_-GspS complex with C5 symmetry applied. A, Both GspD_β_ and GspS are overexpressed; B, Only GspD_β_ is overexpressed. The three units within one C5 symmetry unit are indicated by red arrows.

**Supplementary Figure 7.**
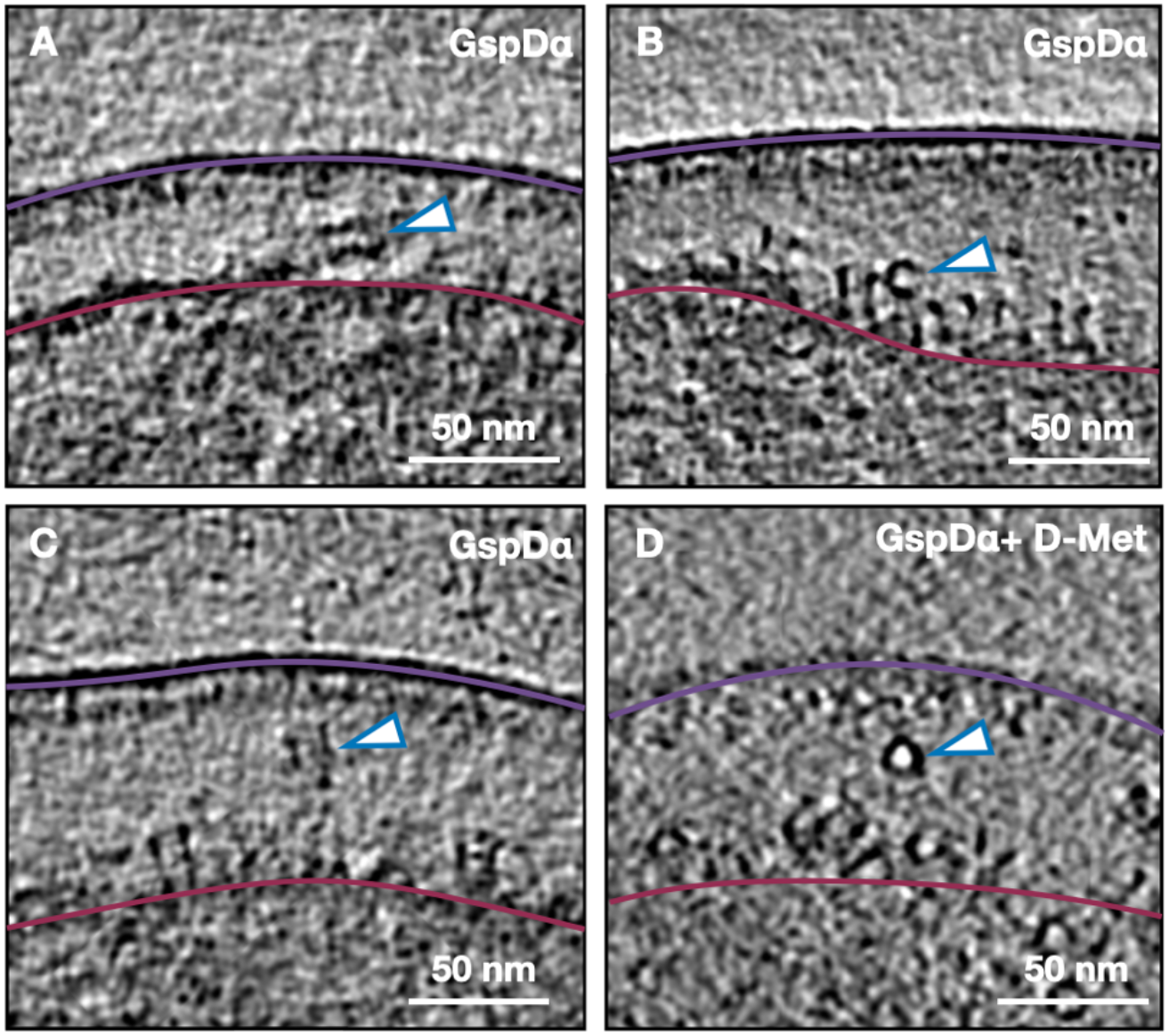
GspD_α_ multimer particles in the periplasm. A, B, and C, Tomogram z slice view of the *E. coli* cells overexpressing GspD_α_. D, Tomogram z slice view of an *E. coli* cell overexpressing GspD_α_ and with D-methionine added. The outer and inner membranes are indicated by the purple line and the dark-red line, respectively. The particles inside the periplasm are indicated by white arrowheads with blue outlines. D-Met, D-methionine.

**Supplementary Figure 8.**
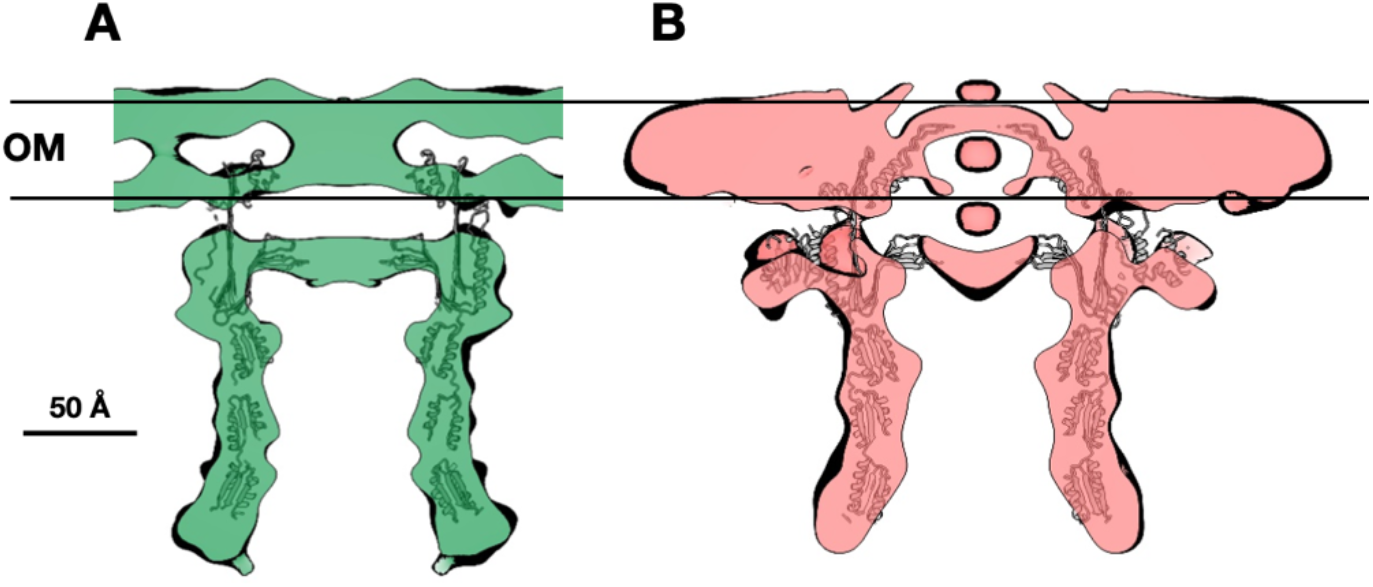
Comparison of the transmembrane regions of GspD_α_ and GspD_β_ *in situ* structures on the outer membrane. A, GspD_α_ on the outer membrane. B, GspD_β_ on the outer membrane. OM, outer membrane.

## Supplementary Movies

**Supplementary Movie 1. Visualization of GspD**_**α**_ **on the *E. coli* inner membrane**. IM, inner membrane; OM, outer membrane; PG, peptidoglycan.

**Supplementary Movie 2. The symmetry release of GspD**_**α**_ **and the movement of GspD**_**α**_ **on the inner membrane**. IM, inner membrane.

## Methods

### Bacterial strains, plasmid construction, and protein expression

*E. coli* BW25113-Δ*gspS* strain was obtained from the Nara Institute of Science and Technology in Japan, from the Keio collection^28,29^. *E. coli* BL21 (DE3)-Δ*gspS* strain was constructed by Ubigene Biosciences Co., Ltd (Guangzhou, China). The DNA sequences encoding GspD_α_ (*E. coli* K12 strain), GspD_β_ (ETEC H10407 strain), and GspS (ETEC H10407 strain) were synthesized by General Biosystems Co., Ltd (Anhui, China).

The *gspD*_*α*_ from the *E. coli* K12 strain with no tag or with a C-terminal hexahistidine tag was cloned into pETDuet-1, yielding pETDuet-*gspD*_*α*_ and pETDuet-*gspD*_*α*_-his, which were expressed in *E. coli* BL21 (DE3), respectively. The vector with no tag was used for cryo-ET imaging, and the vector with the hexahistidine tag was used to perform biochemistry tests. The *gspD*_*β*_ of ETEC H10407 strain with hexahistidine tag on the C terminal was inserted into pETDuet-1, yielding pETDuet-*gspDβ*-his, which was expressed in *E. coli* Rosetta (DE3) and used for cryo-ET imaging and expressed in *E. coli* BL21 (DE3) for biochemistry experiments. The gene of *gspS* was cloned into pETDuet-*gspDβ*-his with a hexahistidine tag at the C terminus, yielding pETDuet-*gspDβ*-his-*gspS*-his, which was expressed in *E. coli* BL21 (DE3) and used for the biochemistry experiments. The *gspD*_*β*_ with hexahistidine tag and *gspS* with no tag from ETEC H10407 strain were both inserted into pETDuet-1’s two multi-cloning sites, yielding pETDuet-*gspDβ*-his-*gspS*, which was expressed in *E. coli* BL21 (DE3) and used for cryo-ET imaging. The *gspD*_*β*_ of ETEC H10407 strain with hexahistidine tag was inserted into pBAD, yielding pBAD-*gspDβ*-his, which was expressed in BW25113-Δ*gspS* strain and used for cryo-ET imaging, and was expressed in BL21 (DE3)-Δ*gspS* strain and used for biochemistry experiments. To obtain minicells expressing the protein of interest, *E. coli* strains were co-transformed with pBS58, which increases the frequency of cell division by constitutively expressing cell division-related genes^30^. The obtained minicell thickness ranges from 130 nm to 250 nm. *E. coli* cells only transformed with pBS58 and not with any protein expression vectors were used as negative control cells.

Cells were grown at 37°C in LB medium supplemented with appropriate antibiotics (final concentrations of 100 *µ*g/ml ampicillin and/or 50 *µ*g/ml kanamycin) and protein expression was induced at an OD_600_ of 0.8, by adding 0.5 mM IPTG or 0.02% arabinose at 20°C overnight. Protein expression was examined by western blotting analysis using anti-His tag antibodies.

### Sample preparation

To perform minicell separation, *E. coli* cultures were first centrifuged at 1,000 × *g* for 40 min to precipitate large cells. The supernatant was taken and centrifuged at 20,000 × *g*, for 10 min. The precipitation was washed with PBS buffer for one time, then resuspended to an OD_600_ of 10. Cells were mixed with 6 nm BSA fiducial gold (Aurion) and then, 3 *µ*l mixture was deposited onto freshly glow-discharged, continuous carbon film-covered grids (Quantifoil Cu R3.5/1 with 2 nm continuous carbon film, 200 mesh). Using Vitrobot Mark IV (FEI), the grids were blotted with filter papers for 4 s and plunged into liquid ethane. Samples were stored in liquid nitrogen.

### Cryo-ET data collection

All tilt series were collected using 5 degrees angular step, from -50 to +50 degrees, defocus from -1.5 *µ*m to -5 *µ*m, and a total dose of about 100 e^−^/Å^2^. For the GspD_α_ on the inner and outer membrane dataset, the sample was imaged using the bidirectional data collection scheme (starting angle -30 degrees) on a 300 kV FEI Titan Krios microscope with a Gatan K2 Summit direct electron detector camera. The magnification was 81,000 ×, with a pixel size of 1.76 Å. For the other datasets and the control dataset, the sample was imaged using a dose symmetric data collection scheme on a 200 kV FEI Glacios microscope with a Falcon 4 direct electron detector camera. The magnification was 92,000 ×, with a calibrated pixel size of 1.52 Å (Supplementary Table 1).

### Cryo-ET data processing of GspD_α_ on the inner membrane dataset EMAN2 refinement pipeline

The raw frames were aligned by MotionCorr2. The tilt series alignment, tomogram reconstruction, and CTF correction were performed in EMAN2^23^. For the refinement, 32,915 particles were manually picked from 250 tilt series and extracted using a box size of 256. After generating an initial model with a small set of particles that have a good signal-to-noise ratio, 4 iterations of subtomogram refinement were performed. The worst 20% of particles were excluded based on their similarity to the averaged structure.

### Using the geometric aspect of the cell membrane to assist particle orientation determination

Due to the shape of GspD_α_, its top view and bottom view are hard to distinguish, and after the refinement, many top view particles were wrongly aligned by about 180 degrees. To correct this, an algorithm was developed that decides a center for each tomogram based on all the particles picked in this tomogram and draws a vector from the particle position to the center (or the opposite direction if the parameter “invert” is toggled). If a particle’s orientation from the refinement result is not facing the same side as its corresponding vector, its orientation will be rotated 180 degrees. After the correction, a new particle orientation file was written and used as input for the next iteration of refinement. In the next iterations, local refinement was used to make sure that the particle orientation will not rotate back to the wrong side. And 3 more iterations of refinement were performed to achieve the final structure.

### Symmetry determination and particle radius measurement

To determine the particle symmetry, we first erected each particle based on its orientation from the refinement and projected them to 2D images, and then performed 2D classification in EMAN2; but the result did not show distinctly different classes or explicit symmetry. We also did refinement with C12, C14, and C16 symmetry, but the resulting density map did not show any obvious feature differences or feature improvements compared to the C15 structure (data not shown). To measure the particle radius, we applied a low pass filter to the particle 2D projection image, calculated the mean radial intensity along the radial axis, and then recorded the index corresponding to the maximum intensity value (Supplementary Figure 2). The particle radius is pixel number times angstrom per pixel.

### Symmetry release and particle movement trajectory calculation

To visualize the membrane connecting region of GspD_α_ more clearly, we did a focused classification, using a mask that is focusing on the membrane connecting region of the density map (Supplementary Figure 3, dashed line box). Within the results, there is one class that shows C1 features (Supplementary Figure 3, red line box). To achieve this C1 structure, we did a symmetry release using the whole dataset: with the orientation of the symmetry axis unaltered, each particle is allowed to rotate around the symmetry axis and search for a best-fitted symmetry unit for one iteration, followed by averaging of subtomograms. Then, the particle orientations are subject to 2 iterations of gold standard refinement, doing only a local search.

We have observed tilted inserted particles from the tomograms (Figure 1E-G), which indicates that, for these particles in the alignment process, if we give a reference map where the membrane and the particle symmetry axis are perpendicular, the membrane density and the protein density could not be aligned correctly at the same time. If the membrane density plane is correctly aligned, the protein density orientation will be inaccurate. To resolve this problem, we did a focused refinement. A mask was generated enclosing the non-transmembrane part of the protein, excluding the membrane density, and one iteration of alignment was performed using the masked structure as a reference (Supplementary Figure 3G). In this iteration, the protein density part should be aligned correctly. We then compared the orientation from the focused refinement with that from the overall refinement, and the orientation difference represents the protein tilt angle. The orientation difference for each particle could be plotted, and a trajectory was calculated. To show the movement, the full trajectory was sectioned into several intervals, and class averages for each interval were calculated representing the corresponding conformation (Supplementary Figure 3H).

### Cryo-ET data processing of the other datasets

The data processing workflow for the GspD_α_ on the outer membrane dataset, GspD_β_-GspS complex dataset, and GspD_β_ on the inner membrane dataset resembles the GspD_α_ on the inner membrane dataset, but simpler. After motion correction and tomogram reconstruction, particles were picked from tomograms. After generating the initial model, the refinement was firstly done with C15 symmetry for 4 iterations, and then particle orientations were corrected using the cell membrane geometry. Then another 4 iterations of local refinement were done to achieve the final structure (for C5 refinements, we only changed the symmetry input to C5, and other parameters were the same). The symmetry release protocol of the GspD_α_ on the outer membrane dataset followed that of the GspD_α_ on the inner membrane dataset.

### Extraction of outer membrane protein

Outer membrane proteins were extracted using a bacterial membrane protein extraction kit (BestBio). Briefly, *E. coli* cells expressing the protein of interest were harvested by centrifugation, washed twice with PBS, and suspended in Extracting Solution A (containing protease inhibitor mixture). After 1 h of shaking at 4°C, the suspension was centrifuged at 12,000 × *g* for 5 min at 4°C to remove the pellet. The supernatant was taken and incubated at 37°C for 1 h, after which it could be observed that the liquid was divided into two layers. The liquid at the bottom was collected as the outer membrane proteins.

#### Separation of the *E. coli* envelope into inner and outer membrane fractions

Separation of inner and outer membranes by isopycnic sucrose density gradient centrifugation was performed as described previously^31^. Briefly, *E. coli* cells expressing the protein of interest were harvested by centrifugation and suspended in buffer A (10 mM Tris-HCl, pH 7.5; 0.5 M sucrose; 10 mg/ml lysozyme; 1.5 mM EDTA). After incubation on ice for 7 min, the suspension was centrifugated at 10,000 × *g* for 10 min at 4°C, and then the pellet was resuspended in buffer B (10 mM Tris-HCl, pH 7.5; 0.2 M sucrose; 1 M MgCl_2_) containing RNase/DNase nuclease reagent and protease inhibitor cocktail. The mixture was lysed by sonication and centrifuged at 6,169 × *g* for 10 min at 4°C to remove the cell debris. Subsequently, the supernatant was pelleted by ultracentrifugation at 184,500 × *g* for 1 h at 4°C and then suspended in buffer C (1 mM Tris-HCl, pH 7.5; 1 mM EDTA,) containing 20% sucrose to get the total membrane fraction. After that, buffer C containing 73% sucrose and buffer C containing 53% sucrose were layered in a 13 ml tube (Beckman Coulter), starting from the bottom, and finally, 0.5 ml of the total membrane fraction was layered on the top, followed by buffer C containing 20% sucrose to fill up the 13 ml tube. Then, the density gradients were centrifuged at 288,000 × *g* for 16 h at 4°C, and the tawny band at the interface between 20% and 53% sucrose layers and the white band at the interface between 53% and 73% sucrose layers were collected as inner membrane fraction and outer membrane fraction, respectively.

## Data and materials availability

Density maps are available at EMDB with accession codes xxx. Other data supporting this study is available from the corresponding author upon reasonable request.

## Acknowledgments

This work was supported by R01GM143380, R01HL162842, and BCM BMB department seed funds to Z.W.; the National Natural Science Foundation of China (No. 82072312), and the Natural Science Foundation of Jiangsu Province (No. BK20211053) to X.S.; Postgraduate Research & Practice Innovation Program of Jiangsu Province (No. YCX22_2925); and NIH R01GM080139 to S.J.L. CryoEM data was collected at the Baylor College of Medicine CryoEM ATC, which includes equipment purchased under the support of CPRIT Core Facility Award RP190602. We thank Dr. Jun Liu and Dr. Bo Hu for sharing the pBS58 plasmid. We thank Hongjiang Wu for discussions about vector design and biochemistry experiments, Valerie Dalton for revising the manuscript language, and Snekalatha Raveendran for data backup.

## Author contributions

Z.W., X.S., and Z.Y. designed experiments. Z.Y. performed sample freezing and cryo-ET imaging. Z.Y., M.C., and T.H. did cryo-ET data processing. Y.W. and W.Z. performed biochemistry experiments. Z.Y. wrote the initial manuscript. Z.W., X.S., S.J.L., and M.C. revised the manuscript.

## References

1. Costa, T. R. D. et al. Secretion systems in Gram-negative bacteria: structural and mechanistic insights. Nat Rev Microbiol 13, nrmicro3456 (2015).

2. Korotkov, K. V., Sandkvist, M. & Hol, W. G. J. The type II secretion system: biogenesis, molecular architecture and mechanism. Nat Rev Microbiol 10, 336 (2012).

3. Cianciotto, N. P. & White, R. C. Expanding Role of Type II Secretion in Bacterial Pathogenesis and Beyond. Infect Immun 85, e00014–17 (2017).

4. Dunstan, R. A. et al. Assembly of the Type II Secretion System such as Found in Vibrio cholerae Depends on the Novel Pilotin AspS. Plos Pathog 9, e1003117 (2013).

5. Yan, Z., Yin, M., Xu, D., Zhu, Y. & Li, X. Structural insights into the secretin translocation channel in the type II secretion system. Nat Struct Mol Biol 24, 177–183 (2017).

6. Hay, I. D., Belousoff, M. J. & Lithgow, T. Structural Basis of Type 2 Secretion System Engagement between the Inner and Outer Bacterial Membranes. Mbio 8, e01344–17 (2017).

7. Hay, I. D., Belousoff, M. J., Dunstan, R. A., Bamert, R. S. & Lithgow, T. Structure and Membrane Topography of the Vibrio-Type Secretin Complex from the Type 2 Secretion System of Enteropathogenic Escherichia coli. J Bacteriol 200, e00521–17 (2018).

8. Yin, M., Yan, Z. & Li, X. Structural insight into the assembly of the type II secretion system pilotin–secretin complex from enterotoxigenic Escherichia coli. Nat Microbiol 3, 581–587 (2018).

9. Chernyatina, A. A. & Low, H. H. Core architecture of a bacterial type II secretion system. Biorxiv 397794 (2018) doi:10.1101/397794.

10. Howard, S. P. et al. Structure and assembly of pilotin-dependent and -independent secretins of the type II secretion system. Plos Pathog 15, e1007731 (2019).

11. Vanderlinde, E. M., Strozen, T. G., Hernández, S. B., Cava, F. & Howard, S. P. Alterations in Peptidoglycan Cross-Linking Suppress the Secretin Assembly Defect Caused by Mutation of GspA in the Type II Secretion System. J Bacteriol 199, e00617–16 (2017).

12. Vanderlinde, E. M. et al. Assembly of the Type Two Secretion System in Aeromonas hydrophila Involves Direct Interaction between the Periplasmic Domains of the Assembly Factor ExeB and the Secretin ExeD. Plos One 9, e102038 (2014).

13. Zhang, S. et al. Scaffolding Protein GspB/OutB Facilitates Assembly of the Dickeya dadantii Type 2 Secretion System by Anchoring the Outer Membrane Secretin Pore to the Inner Membrane and to the Peptidoglycan Cell Wall. Mbio 13, e00253–22 (2022).

14. Strozen, T. G., Li, G. & Howard, S. P. YghG (GspSβ) Is a Novel Pilot Protein Required for Localization of the GspSβ Type II Secretion System Secretin of Enterotoxigenic Escherichia coli. Infect Immun 80, 2608–2622 (2012).

15. Konovalova, A. & Silhavy, T. J. Outer membrane lipoprotein biogenesis: Lol is not the end. Philosophical Transactions Royal Soc B Biological Sci 370, 20150030 (2015).

16. Francetic, O. & Pugsley, A. P. The cryptic general secretory pathway (gsp) operon of Escherichia coli K-12 encodes functional proteins. J Bacteriol 178, 3544–3549 (1996).

17. Francetic, O., Belin, D., Badaut, C. & Pugsley, A. P. Expression of the endogenous type II secretion pathway in Escherichia coli leads to chitinase secretion. Embo J 19, 6697–6703 (2000).

18. d’Enfert, C., Reyss, I., Wandersman, C. & Pugsley, A. P. Protein secretion by gram-negative bacteria. Characterization of two membrane proteins required for pullulanase secretion by Escherichia coli K-12. J Biological Chem 264, 17462–8 (1989).

19. d’Enfert, C., Ryter, A. & Pugsley, A. P. Cloning and expression in Escherichia coli of the Klebsiella pneumoniae genes for production, surface localization and secretion of the lipoprotein pullulanase. Embo J 6, 3531–3538 (1987).

20. Horstman, A. L. & Kuehn, M. J. Bacterial Surface Association of Heat-labile Enterotoxin through Lipopolysaccharide after Secretion via the General Secretory Pathway. J Biol Chem 277, 32538–32545 (2002).

21. Yang, J., Baldi, D. L., Tauschek, M., Strugnell, R. A. & Robins-Browne, R. M. Transcriptional Regulation of the yghJ-pppA-yghG-gspCDEFGHIJKLM Cluster, Encoding the Type II Secretion Pathway in Enterotoxigenic Escherichia coli. J Bacteriol 189, 142–150 (2007).

22. Farley, M. M., Hu, B., Margolin, W. & Liu, J. Minicells, Back in Fashion. Journal of Bacteriology 198, 1186–1195 (2016).

23. Chen, M. et al. A complete data processing workflow for cryo-ET and subtomogram averaging. Nat Methods 16, 1161–1168 (2019).

24. Korotkov, K. V., Delarosa, J. R. & Hol, W. G. J. A dodecameric ring-like structure of the N0 domain of the type II secretin from enterotoxigenic Escherichia coli. J Struct Biol 183, 354–362 (2013).

25. Reichow, S. L., Korotkov, K. V., Hol, W. G. J. & Gonen, T. Structure of the cholera toxin secretion channel in its closed state. Nat Struct Mol Biol 17, 1226 (2010).

26. Caparrós, M., Pisabarro, A. G. & Pedro, M. A. de. Effect of D-amino acids on structure and synthesis of peptidoglycan in Escherichia coli. J Bacteriol 174, 5549–5559 (1992).

27. Huang, K. C., Mukhopadhyay, R., Wen, B., Gitai, Z. & Wingreen, N. S. Cell shape and cell-wall organization in Gram-negative bacteria. Proc National Acad Sci 105, 19282–19287 (2008).

28. Baba, T. et al. Construction of Escherichia coli K-12 in-frame, single-gene knockout mutants: the Keio collection. Mol Syst Biol 2, 2006.0008-2006.0008 (2006).

29. Yamamoto, N. et al. Update on the Keio collection of Escherichia coli single-gene deletion mutants. Mol Syst Biol 5, 335–335 (2009).

30. Hu, B. et al. Visualization of the type III secretion sorting platform of Shigella flexneri. Proceedings of the National Academy of Sciences 112, (2015).

31. Cian, M. B., Giordano, N. P., Mettlach, J. A., Minor, K. E. & Dalebroux, Z. D. Separation of the Cell Envelope for Gram-negative Bacteria into Inner and Outer Membrane Fractions with Technical Adjustments for Acinetobacter baumannii. J Vis Exp (2020) doi:10.3791/60517.

